# Ethanol-guided behavior in *Drosophila* larvae

**DOI:** 10.1101/2021.02.07.430116

**Authors:** Isabell Schumann, Michael Berger, Nadine Nowag, Yannick Schäfer, Juliane Saumweber, Henrike Scholz, Andreas S. Thum

## Abstract

Chemosensory signals allow vertebrates and invertebrates not only to orient in its environment toward energy-rich food sources to maintain nutrition but also to avoid unpleasant or even poisonous substrates. Ethanol is a substance found in the natural environment of *Drosophila melanogaster*. Accordingly, *D. melanogaster* has evolved specific sensory systems, physiological adaptations, and associated behaviors at its larval and adult stage to perceive and process ethanol.

To systematically analyze how *D. melanogaster* larvae respond to naturally occurring ethanol, we examined ethanol-induced behavior in great detail by parametrically reevaluating existing approaches and comparing them with new experiments. Using behavioral assays, we confirm that larvae are attracted to different concentrations of ethanol in their environment. This behavior is controlled both by olfactory and contact cues. It is independent of previous exposure to ethanol in their food. Moreover, moderate, naturally occurring ethanol concentration of 4% results in increased larval fitness. On the contrary, higher concentrations of 10% and 20% ethanol, which rarely or never appear in nature, increase larval mortality. Finally, ethanol also serves as a positive teaching signal in learning and memory and updates valence associated with simultaneously processed odor information.

Since information on how larvae perceive and process ethanol at the genetic and neuronal level is limited, the establishment of standardized assays described here is an important step towards their discovery.

## Introduction

Communication with the environment through chemical signals is an essential process for the survival of most if not all organisms. Specialized signal transduction pathways are used to detect chemical cues and convert information into neuronal activity that induces appropriate behavioral output, which is evolutionary conserved between invertebrates and vertebrates, identified by biochemical, molecular and genetic approaches^1^. Important insights into principals of chemosensory perception and information processing are provided by genetically modifiable organisms such as the fruit fly *D. melanogaster* ^2–5^. This includes also the larval central nervous system with its simpler structure consisting of only about 10,000 neurons, but equally provides access to combinations of genetic tools, robust behavioral assays, the possibility of transgenic single-cell manipulation, and even connectome data of the central nervous system ^6–14^. Various studies have identified chemosensory stimuli that larvae perceive from their environment. Most odorants, are attractive to larvae in a dose-dependent manner ^15–17^. Likewise, larvae show dose-dependent responses to gustatory cues ^18–22^. Even the characteristics of the substrate seem to influence larval chemosensory responses ^23^,^24^. However, to understand how larvae orient in their complex chemosensory environment, further studies are needed on additional environmental occurring stimuli, such as ethanol.

Most *Drosophila* species are saprophagous and feed on decaying sweet substrates like rotting fruits, which contain ethanol produced by natural fermentation ^25^. Adult flies exhibit acute ethanol responses similar to those of mammals: as ethanol concentration increases, flies exhibit locomotor stimulation, loss of postural control, and eventually sedation ^26^. With repeated exposure, adults develop tolerance to the effects of ethanol ^27^. It is even assumed that *D. melanogaster* has altered its ecological niche to benefit from food sources characterized by a higher alcohol concentration ^28^. *D. melanogaster* has an unusually high alcohol dehydrogenase (Adh) activity within its genus, which allows to deal with higher ethanol levels to occupy microhabitats that are not accessible to other species such as *Drosophila simulans*, which have a lower Adh activity ^29^. In addition, larvae are even able to perceive and selectively consume food containing ethanol when they are infected by parasitic wasps ^30^. A higher ethanol in the larval hemocoel due to increased consumption of ethanol enriched food leads to enhanced death of the growing parasites and thus increases the larval survival rate. This means that *D. melanogaster* larvae can adjust their alcohol consumption depending on the particular situation, and use it as a kind of medical treatment. Accordingly, ethanol is an important ecological parameter, which has selected to the establishment of specific larval sensory systems, physiological adaptations and related behaviors.

Therefore, it is plausible that several studies have been able to show that *D. melanogaster* larvae preferentially migrate to substrates that contain ethanol. For instance, different wild-type strains derived from Australian populations revealed strong preferences for 6% ethanol ^31–33^. Even 17% ethanol was attractive for two larval strains that were either homozygous for the Adh^F^ or the Adh^S^ allele ^34^. Adh^F^ and Adh^S^ describe two functional allozymes that exist in natural populations of *Drosophila* species on several continents ^35^,^36^. The polymorphism is maintained by natural selection; while Adh^F^ is showing a higher enzymatic activity for ethanol, Adh^S^ is more resistant to higher environmental temperatures ^37^. However, there are also studies in which ethanol is not attractive to larvae ^15^,^38^,^39^, which complicates the interpretation of the various results. Therefore, we parametrically analyzed ethanol guided behavior by combining published and novel experimental designs.

In recent years, a set of well-defined larval behavioral assays was established to investigate substrate choice, feeding, survival, and associative learning and memory ^14^,^18^,^19^. It is therefore possible to analyze how larvae react to individual chemical components of its environment. Fructose, for example, provides nutrition and acts as a positive teaching signal during learning ^20^,^21^. On the other hand, many chemicals that humans categorize as bitter are also repulsive to the larvae (for example among others caffeine, denatonium, or quinine) and can act as a negative teaching signal (caffeine, quinine) ^18^,^19^,^22^,^40^. Sodium chloride even proved to be dichotomous, being attractive and a positive teaching signal at low concentrations and repellent and a negative teaching signal at high concentrations ^41^,^42^. Using a set of standardized assays, we have now addressed the effect of ethanol on larval behavior. Our results support previous studies suggesting that the odorant ethanol is attractive to larvae in a doses dependent manner at moderate naturally occurring concentrations and has a positive effect on larval survival. The larval ethanol choice is based on smell and taste. Finally, ethanol can be used in associative olfactory learning paradigms to establish appetitive memories as it provides a positive teaching signals.

## Results

### *Drosophila melanogaster* larvae are attracted to ethanol

Several studies have repeatedly shown that *D. melanogaster* larvae are attracted to ethanol ^31–34^. However, in almost all studies there are slight differences in the applied assays, used substrates and mode of ethanol presentation. This might influence the observed results. In some cases ethanol is attractive and in other not ^15^,^38^,^39^. Therefore, we first reanalyzed the attractiveness of ethanol in a widely applied standardized assay. We analyzed the preference of wild-type *Canton-S* to ethanol by observing their approach behavior to a substrate that contains different concentration of ethanol (Fig. 1). The ethanol concentration ranged from 1 % to 30 %. We found that *Canton-S* and *w^1118^* larvae are attracted to ethanol in a dose-dependent manner (Fig. 1b). The attractiveness of ethanol peaked at around 4 – 10% ethanol for *Canton-S*. For 10% and 20% ethanol *Canton-S*-larvae showed higher preferences than *w^1118^* larvae. Additionally, we observed the attractiveness to 8% ethanol of the *Canton-S* larvae for 120 min (Fig. 1c). The attraction to ethanol remained equally stable and no sedative effect on larval locomotion was seen during this prolonged ethanol exposure. To investigate the influence of ethanol pre-exposure on the substrate choice of larvae we compared how animals reared on standard food (containing 1% ethanol) versus animals reared for one or two generations on ethanol-free food respond to 8% ethanol (Fig. 1d). *Canton-S* larvae grown for one or two generations on ethanol-free food preferred the ethanol containing site similar to larvae raised on standard food. These results show that *D. melanogaster* larvae are attracted to ethanol in a concentration-dependent manner. Pre-exposure to ethanol and the duration of the test period do not alter the attraction.

**Figure 1:**
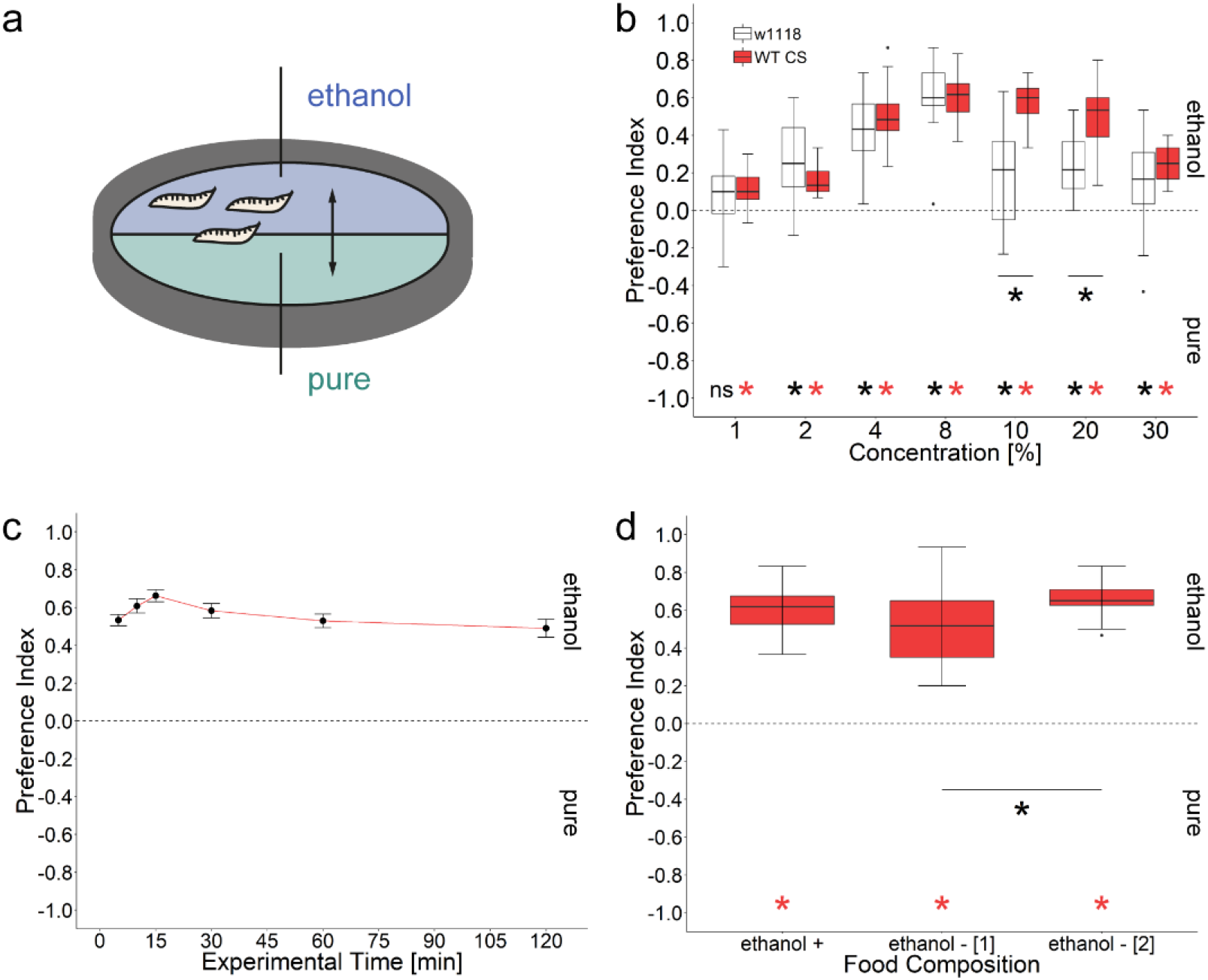
Ethanol attraction of *Drosophila melanogaster* larvae. (a) Scheme of the experimental procedure. Larvae were allowed to choose within five minutes between pure agarose and a substrate containing different concentrations of ethanol. (b) Wild-type *Canton-S* (red) and mutant *w^1118^* (white) larvae are attracted to ethanol. The highest behavioral response (preference index (*Pref*)) was seen at 8% ethanol for both strains (*Pref^Canton-S^*= 0.62, ci = 0.53–0.68, *Pref^w1118^*= 0.60, ci = 0.51–0.74). All groups are significant different to zero, except for 1% *w^1118^* (t-test, *p* < 0.17). Multiple comparison indicates a significant difference between the two genotypes for 10% and 20% ethanol (Wilcoxon-Rank, *p^10%^* < 0.001, *p^20%^* < 0.001). (c) Larval 8% ethanol substrate choice over a test period of 120 min. *Canton-S* larvae did not vary in their attraction within 120 min to 8% ethanol (Kruskal Wallis, p < 0.032). (*Pref^5min^* = 0.55, ci = 0.47–0.6, *Pref^10min^* = 0.65, ci = 0.53–0.69, *Pref^15min^* = 0.63, ci = 0.6–0.73, *Pref^30min^* = 0.53, ci = 0.5–0.67, *Pref^60min^* = 0.5, ci = 0.45–0.61, *Pref^120min^* = 0.57, ci = 0.39–0.59). Indicated values show medians; error bars represent standard errors. (d) *Canton-S* larvae raised on standard food that contains 1% ethanol (ethanol +) shows a substrate choice behavior that was not different from larvae that were raised for one (ethanol - [1]) or two generations (ethanol - [2]) on ethanol free food (TukeyHSD, *p^standard-w/o^* < 0.202, *p^standarad-w/o2^* < 0.639). Note a slight increase in ethanol substrate choice with increasing generations of ethanol food free raised flies (TukeyHSD, *p* < 0.03). Differences against zero are indicated in red and black at the bottom of each panel. Sample size for each box plot is n = 16. Significant differences of two groups are specifically indicated with an asterisk. Non-significant results are not indicated. Preference scores and statistical tests underlying the different indices are documented in the supplementary material.

### Ethanol attraction changes during post-embryonic development

The life cycle of *D. melanogaster* comprises three larval stages (L1, L2 and L3), clearly separated by two molting stages, which occur approximately 48 and 72 h after egg laying (AEL). To understand whether ethanol choice changes during larval development, we analyzed eight groups of *Canton-S* larvae from 36 – 120 h AEL every 12 h for their 8% ethanol attraction (Fig. 2). We chose this concentration because it was most preferred by the larvae in the first experiment (Fig. 1). Each of the tested groups showed an attraction to ethanol (Tab. 1). However, during the first and second molt, the preferences are reduced (Fig. 2, Tab. 1). Furthermore, in L3 larvae the preference decreases with age.

**Figure 2:**
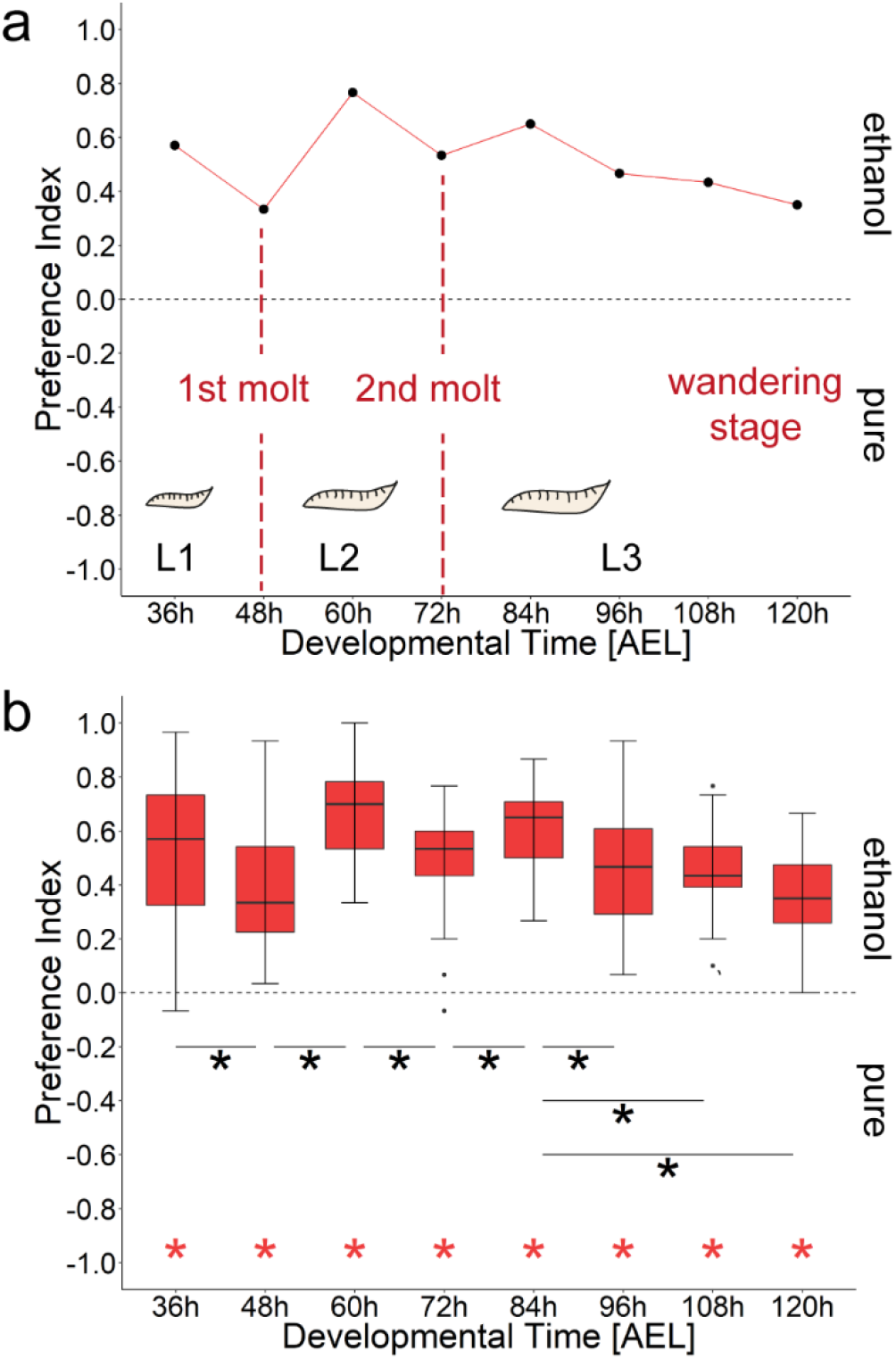
Ethanol substrate choice during larval development. (a) *Canton-S* larvae are attracted to 8% ethanol throughout the post-embryonal development. Data points represent the median behavioral response (substrate choice) according to the time after egg laying (AEL). Note a decrease of the behavioral response to some time points (48h AEL, 72h AEL, 96h–120h AEL) which correlates to larval molting events. (b) Corresponding box plots. Statistics are presented in Table 1. Differences from random choice are indicated in red at the bottom. Significant differences of two groups are specifically indicated with an asterisk. Non-significant results are not indicated.

**Table 1:** Statistical analysis of substrate choice throughout the post-embryonal development of *D. melanogaster*.

| Developmental Time Points (AEL) | N | Preference Index | Statistics<br>Dunn's Multiple Comparison | Event | CI |
| --- | --- | --- | --- | --- | --- |
| 36h | 32 | 0.57 | *<br>$p < 0.001$ | 1st molt | 0.44–0.63 |
| 48h | 24 | 0.33 |  |  | 0.30–0.52 |
| 60h | 31 | 0.77 | *<br>$p < 0.002$ | 2nd molt | 0.59–0.73 |
| 72h | 33 | 0.53 |  |  | 0.41–0.55 |
| 84h | 16 | 0.65 | *<br>$p < 0.001$ | Feeding L3 | 0.51–0.69 |
| 96h | 32 | 0.47 |  |  | 0.38–0.53 |
| 108h | 16 | 0.43 | *<br>Comparison 84h AEL<br>$p < 0.017$ | Wandering L3 | 0.35–0.54 |
| 120h | 16 | 0.35 | *<br>Comparison 84h AEL<br>$p < 0.001$ | | 0.27–0.45 |

### Ethanol has nutritional benefit for the larva

Ethanol can be used as energy source (reviewed in ^43^). However, the nutrient gain for the larva appears to be lower, as survival rates on ethanol substrates are lower than those of adult *Drosophila* ^44,45^ and also than those of larvae on sugar diets ^20^. To complicate matters, ethanol is pharmacologically active and in higher concentrations toxic and needs to be neutralized ^46–48^. These diametrically opposed effects complicate the interpretation of ethanol-dependent survival studies. To resolve and better interpret these effects, we analyzed ethanol dependent survival under standard conditions applied for other environmental cues ^18^,^20^,^23^. We put L2 *Canton-S* larvae in food vials containing 1% agarose an either no ethanol, 4%, 8%, or 20% (Fig. 3). We calculated a survival rate by counting the number of living larvae and pupae for eight days in a 24h cycle (Fig. 3b, c). The results show two effects: at 4% ethanol, more larvae survive than on pure agarose until pupation (73.8% versus 35.8%; Fig. 3b, c, light blue line) and at higher concentration of 20% only few animals survive until pupation (12.2%; Fig. 3b, c, dark blue line). The larval survival rates on pure agarose and 8% ethanol are right in between (Fig. 3b, 44.9% versus 60.9%, green versus blue line). However, of these animals less larvae pupated on 8% ethanol in comparison to pure agarose condition (Fig. 3c).

**Figure 3:**
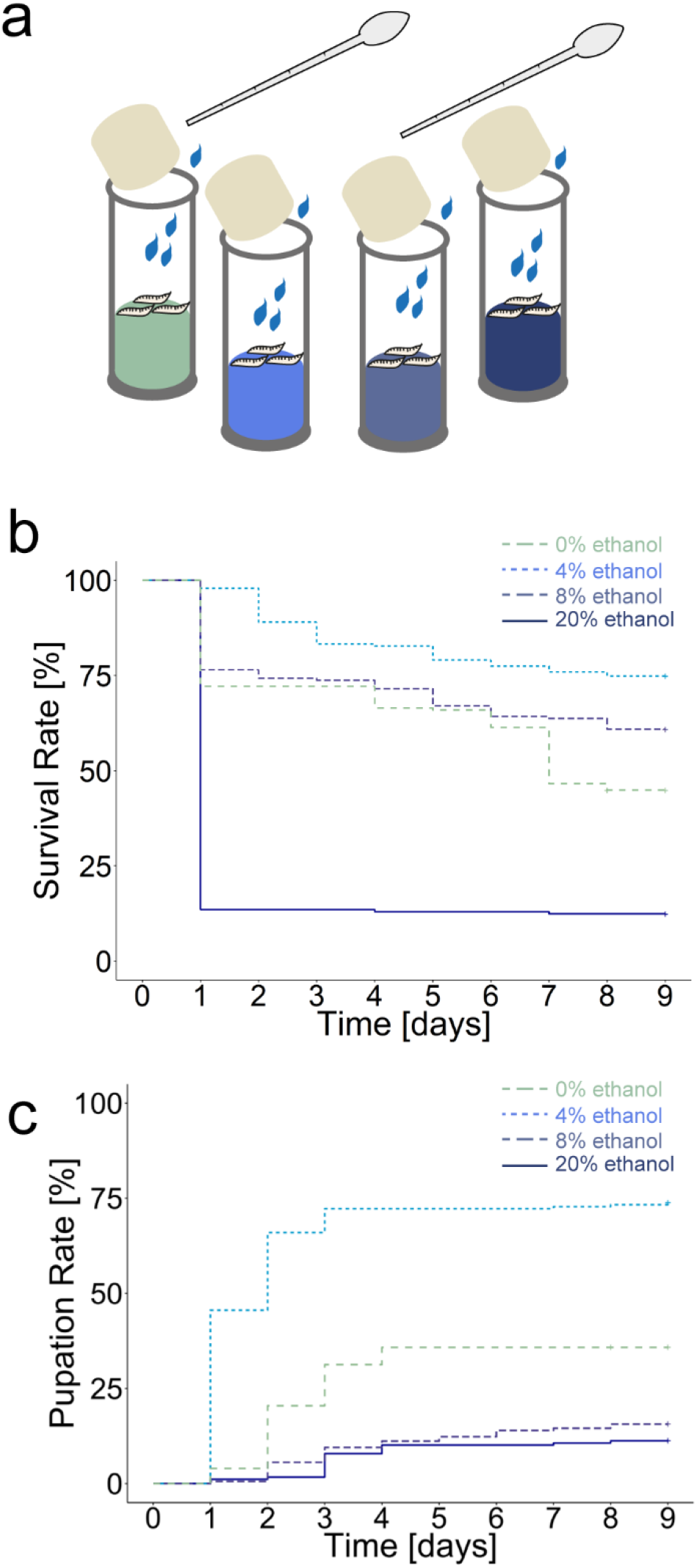
Larval survival on diets containing different concentrations of ethanol. (a) Scheme of the experimental procedure. Independent groups of 12 second instar *Canton-S* larvae were put into food vials that contained an agarose substrate plus different concentration of ethanol. Surviving of larvae and pupae were counted every 24h. Water was added every 24h to avoid dehydration. Pupation was used as measure of survival. (b) Diagram shows Kaplan-Meier survival curves of larvae. Larvae were put either on 0% (green), 4% (light blue), 8% (blue), or 20% ethanol (dark blue) diet for eight days. 20% ethanol significantly reduced larval survival as most of the animals died within one day (*survival rate^20%Ethanol^* = 12.40%). Larvae reared on 0% and 8% ethanol showed a nearly similar survival (Log-Rank-test, p < 0.001, *survival rate^0%Ethanol^* = 44.90%, *survival rate^8%Ethanol^* = 60.90%). In contrast 4% ethanol diet significantly increased larval survival compared to the 0% ethanol control diet (Log-Rank-test, p < 0.001, *survival rate^4%Ethanol^* = 74.90%). (c) Diagram shows the related pupation rates of the surviving animals shown above raised at 0% (green), 4% (light blue), 8% (blue), or 20% ethanol (dark blue) diet. 4% ethanol diet significantly increased the pupation rate compared to larvae reared at 0% ethanol diet (Log-Rank-test, p < 0.001, pupation *rate^0%Ethanol^* = 35.8%, *pupation rate^4%Ethanol^* = 73.80 %). In contrast 8% and 20% ethanol diet significantly reduced the pupation rate compared to larvae reared at 0% ethanol diet (Log-Rank-test, p^0%-8%Ethanol^ < 0.001, p^0%-20%Ethanol^ < 0.001, *pupation rate^8%Ethanol^* = 15.6%, *pupation rate^20%Ethanol^* = 12.2%). Single scores and statistical tests underlying the different survival curves are documented in the supplementary material. Sample size for each Kapplan-Meier survival curve is n = 16.

### Larvae perceive ethanol as an olfactory cue

Initially we performed substrate choice experiments that allowed larvae to get into direct contact with ethanol in addition to smelling the odorant source (Fig. 1, 2). To disentangle the olfactory from contact cues we refined our behavioral approach by presenting ethanol in custom-made Teflon containers with perforated lids. This eliminates a physical contact with the substrate to specifically address the olfactory response of larvae (Fig. 4). We compared the attraction of 8% ethanol to three well-known attractive odorants: 1-octanol (1-OCT), amyl acetate (AM), and benzaldehyde (BA) ^15–17^. *Canton-S* larvae showed olfactory preference for all four odorants (Fig. 4b). Thus, *D. melanogaster* larvae perceive ethanol as an attractive olfactory stimulus. In the next step we analyzed whether the larva can detect other odorants in a homogeneous ethanol background, which is a basic requirement for further odorant-ethanol learning experiments (Fig. 4d). Since BA triggered the greatest behavioral response, we used this stimulus to test whether larvae can still perceive it on 5%, 8%, 10% and 20% ethanol containing agarose plates (Fig. 4d). For all four ethanol concentrations wild-type *Canton-S* larvae were similarly attracted to BA (Fig. 4d). Therefore, in the assay *D. melanogaster* larvae could distinguish BA from a week or high concentrated ethanol background.

**Figure 4:**
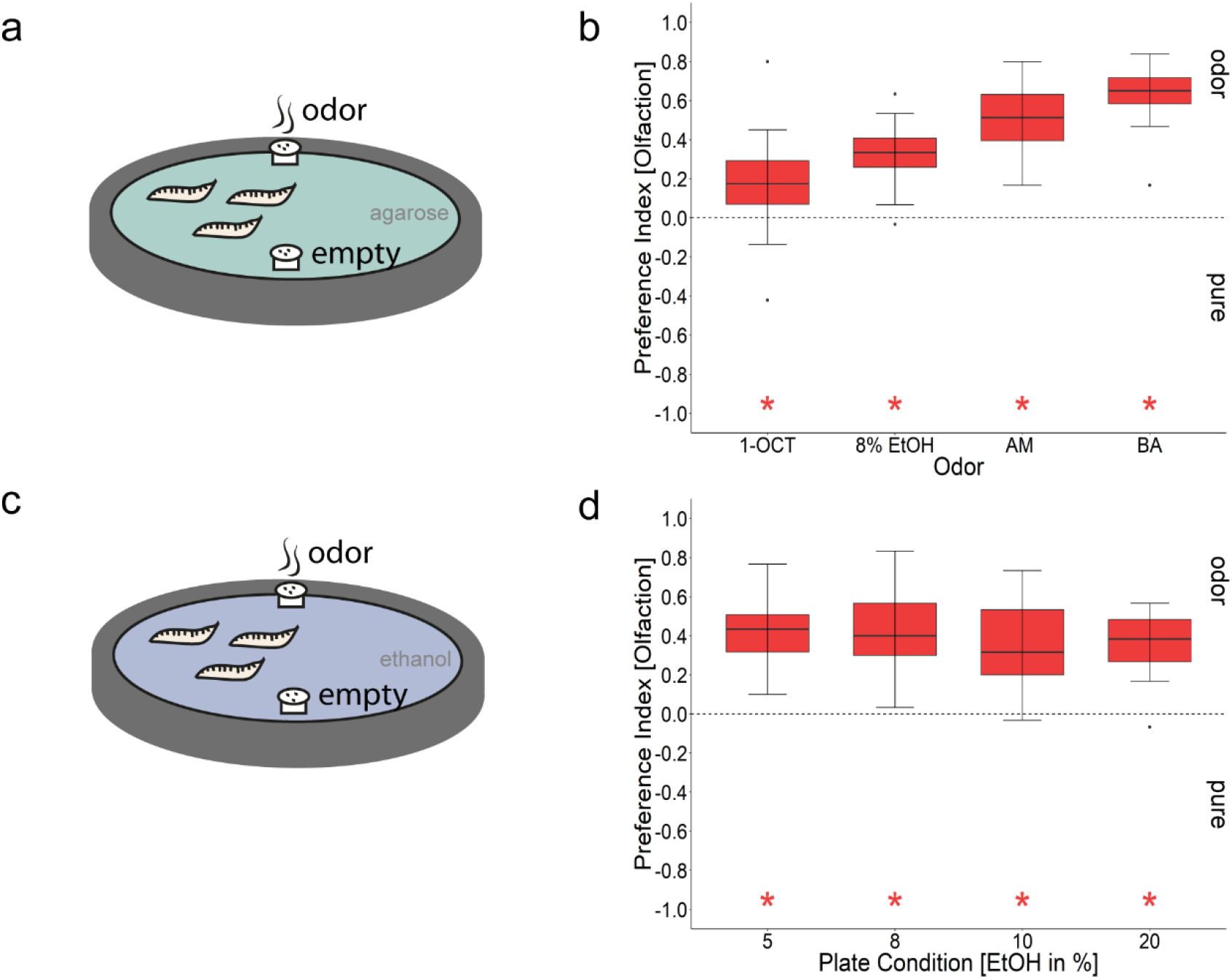
The effects of ethanol on larval olfactory attraction. (a) Scheme of the experimental procedure. The olfactory stimuli 1-OCT, AM, BA, and 8% ethanol were presented on a 2.5% agarose plate in custom made Teflon containers to prevent direct contact of the animals with the chemical. Larvae were allowed to crawl for 5 min. (b) *Canton-S* larvae preferred all odors including 8% ethanol over empty containers (n^1-Octanol^ = 19, *Pref^1-Octanol^* = 0.17, ci = 0.04–0.29, n^AM^ = 8, *Pref^AM^* = 0.51, ci= 0.3–0.69, n^8%Ethanol^ = 16, *Pref^8%Ethanol^* = 0.33, ci = 0.24–0.41, n^BA^ = 16, *Pref^BA^* = 0.65, ci = 0.55–0.72). All groups were significantly different to zero (t-test, *p^1-octanol^* < 0.011, p^AM^ < 0.001, *p^8%Ethanol^* < 0.001, *p^BA^* < 0.001). (c) Scheme of the experimental procedure. Test plates contained 2.5% agarose mixed with either 5%, 8%, 10%, or 20% ethanol. BA was presented in Teflon containers. Larvae were allowed to crawl for 5 min. (d) *Canton-S* larvae showed an olfactory preference for BA irrespective of the ethanol concentrations in the test plate. All groups were significantly different to zero (t-test, *p^BA-5%Ethanol^* < 0.001, *p^BA-8%Ethanol^* < 0.001, *p^BA-10%Ethanol^* < 0.001, *p^BA-20%Ethanol^* < 0.001), but not different from each other (ANOVA, *p* < 0.697). Sample size for each box plot is n = 16. Differences against zero are indicated in red at the bottom of each panel. Significant differences of two groups are specifically indicated with an asterisk. Non-significant results are not indicated. Preference scores and statistical tests underlying the different indices are documented in the supplementary material.

### Ethanol provides a teaching signal for larval olfactory learning

Larvae of *D. melanogaster* are able to associate odorants (conditioned stimulus) with cues of different sensory modalities (unconditioned stimulus) to establish appetitive or aversive olfactory memories. So far, tastants, temperature, vibration, electric shock, and light have been identified as teaching signals (reviewed in ^14^). In contrast to adult flies, ethanol has not yet been tested as reinforcer. Therefore, we performed standardized learning experiments to validate the potential of ethanol in differential conditioning. In this assay larvae are trained three times with a given concentration of ethanol (2.5%, 8%, or 20% as a teaching signal) and then tested on an agarose plate for their odorant preference between the previously ethanol paired and the non-paired odorant (Fig. 5). Higher ethanol concentrations of 8% and 20%, in contrast to a lower concentration of 2.5%, provide an appetitive teaching signal for larval olfactory learning (Fig. 5b). For appetitive olfactory learning with fructose, it has been reported that larvae show no memory when tested in the presence of the unconditioned stimulus. This is probably because in the presence of food the larvae do not have to search for food ^41^,^49^. This also seems to be the case for ethanol, since larvae trained with 8% ethanol only recall odor-ethanol memory on pure agarose, but not on a test plate containing 8% ethanol (Fig. 5c). Ethanol memory also appears to be somehow similar to fructose memory, as even 2M fructose added to the test plate prevents the recall of the odor-ethanol memory.

**Figure 5:**
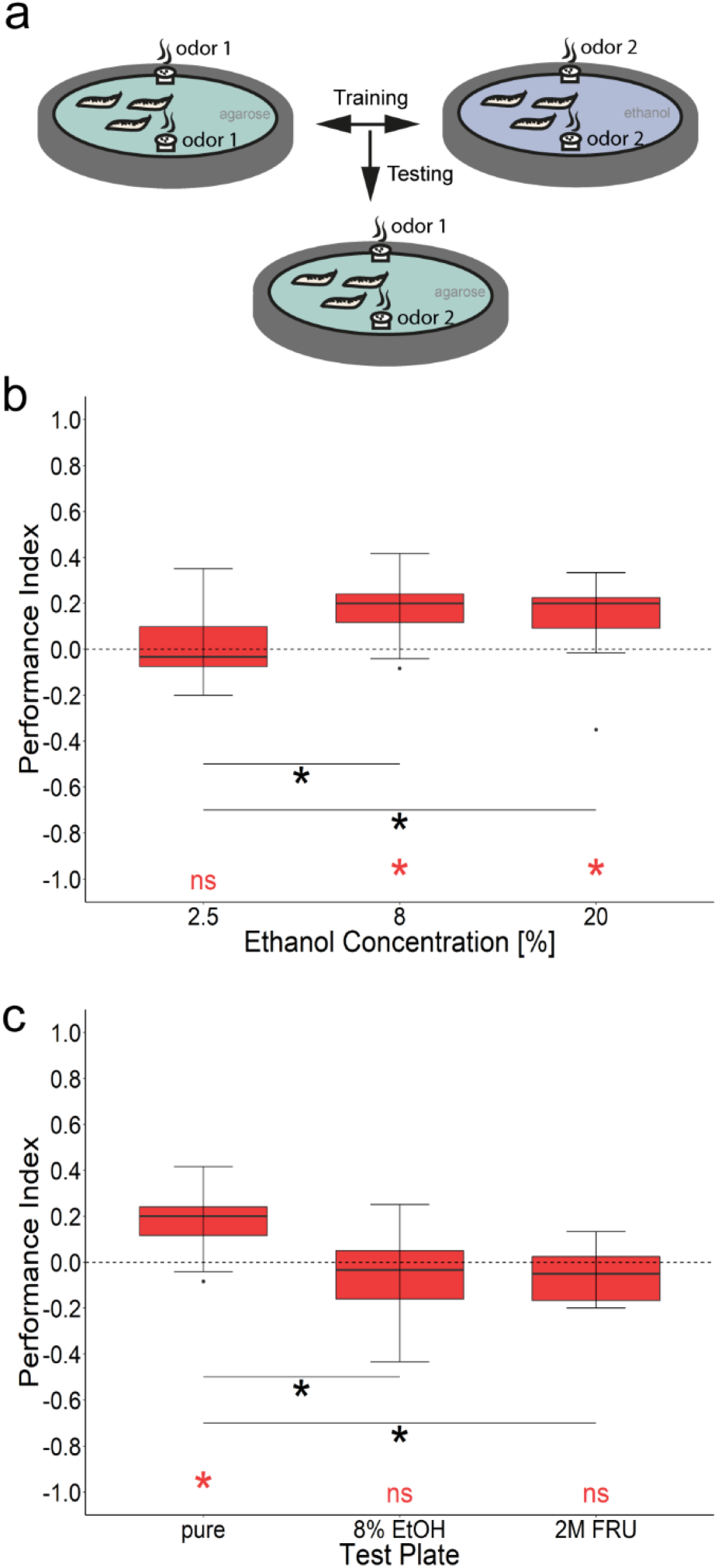
Larval olfactory learning reinforced by three different ethanol concentrations. (a) Scheme of the experimental procedure. *Canton-S* were trained three times with two odorants (AM and BA) and either 2.5%, 8%, or 20% ethanol as reinforcer. Olfactory memory is quantified by the performance index (PI). (b) Larvae trained with 8% and 20% ethanol concentration show a significant appetitive memory (*PI^8%Ethanol^* = 0.20, ci= 0.11–0.26, *PI^20%Ethanol^* = 0.20, ci = 0.05–0.24, t-test, *p^8%^* < 0.001, *p^20%^* < 0.005). When 2.5% ethanol was used as a teaching signal no memory was detectable (*PI^2.5%Ethanol^* = 0.83, ci = −0.07–0.08, t-test, *p^2.5%^* < 0.828). Accordingly, larvae trained with 8% and 20% ethanol behaved differently from larvae trained with 2.5% ethanol (Dunn’s Multiple Comparison test, *p* < 0.001 and *p* < 0.003, respectively). (c) Larvae trained with 8% ethanol concentration as teaching signal and tested on pure agarose showed a significant appetitive memory (*PI* = *0.20*, ci = 0.11–0.26, t-test, p < 0.001); if tested in the presence of the teaching signal of 8% ethanol or 2M fructose (2M FRU), appetitive associative memory is not expressed (*PI^8%Ethanol^* = −0.03, ci = −0.17–0.05, *PI^2MFru^* = −0.05, ci = −0.11–0.01) as it is not significantly different from zero (t-test, *p^8%Ethanol^* < 0.260, *p^2MFru^* < 0.080) and not significantly different from each other (paired t-test, *p* < 0.902). Differences against zero are indicated in red at the bottom of each panel. Sample size for each box plot is n = 15. Significant differences of two groups are specifically indicated with an asterisk. Preference scores and statistical tests underlying the different indices are documented in the supplementary material.

### Ethanol does not alter appetitive and aversive olfactory memory

Treating larvae with 20% ethanol for 20 min impairs aversive odorant-heat shock memory after one-odor conditioning at 35°C ^50^. The effect of ethanol was absent, when the odorant was paired with a higher temperature heat-shock of 41°C, which likely forms a stronger memory unsusceptible for ethanol treatment. Therefore, we similarly investigated whether the 20 min incubation with 20% ethanol interferes with larval olfactory memory formation when the odorant is paired with fructose or salt (Fig. 6). We found that both, aversive salt memory (1.5 M NaCl; Fig. 6a) and appetitive sugar memory (0.01 M fructose; Fig. 6b), showed no reduction after ethanol treatment in comparison to H_2_O treated larvae. Therefore, the olfactory memory reinforced by salt or fructose was not altered by ethanol treatment.

**Figure 6:**
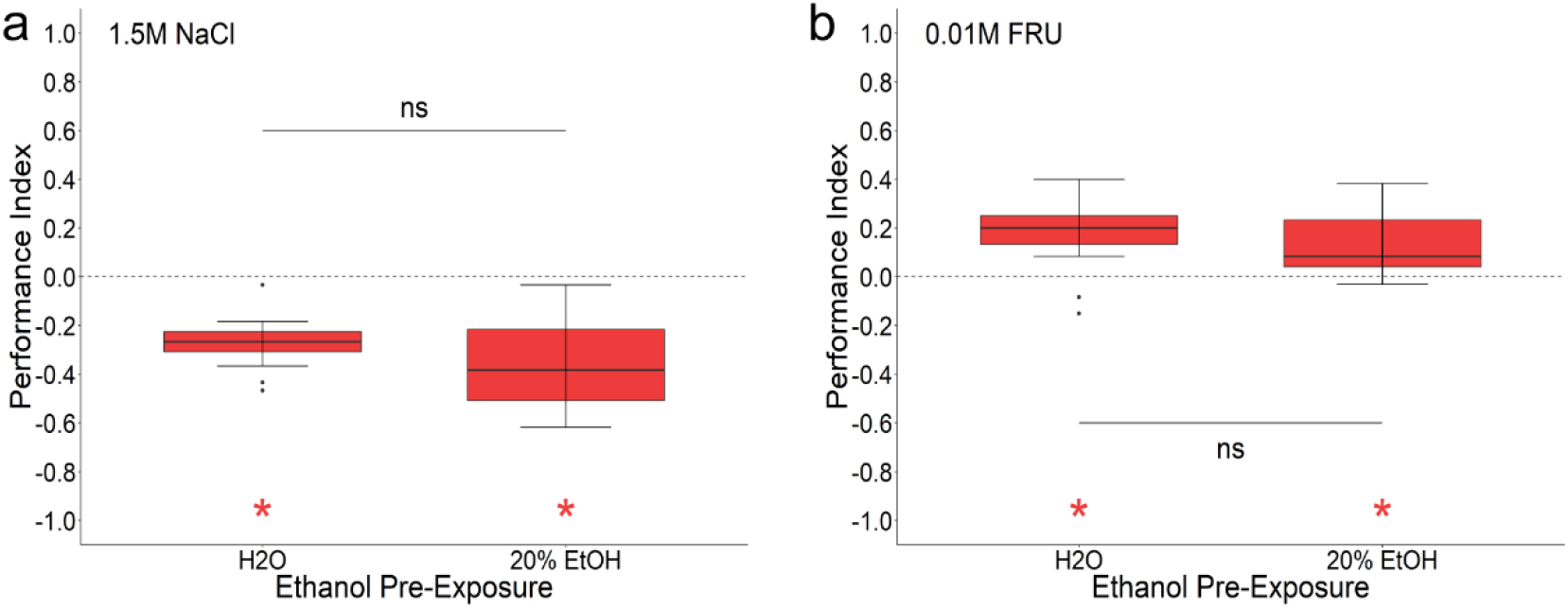
Pre-exposure to ethanol does not change aversive and appetitive olfactory learning. Before the larvae were conditioned, they were separated from their food and treated for 20 minutes with either water or 20% ethanol. They were then washed and trained with the odors AM and BA and either 1.5M NaCl or 0.01M fructose as a teaching signal. (a) Larvae treated with water and 20% ethanol both showed an aversive odor-salt memory (t-test, pwater < 0.001, ppre-EtOH < 0.001) that was not different from each other (paired t-test, p < 0.112). Sample size for each box plot is n = 15. (b) Larvae treated with water and 20% ethanol both showed an appetitive odor-fructose memory (t-test, pwater < 0.001, ppre-EtOH < 0.001) that was not different from each other (paired t-test, p < 0.484). Sample size for each box plot is n = 15. Differences against zero are indicated in red at the bottom of each panel. Significant differences of two groups are specifically indicated with an asterisk. Non-significant results are not indicated. Preference scores and statistical tests underlying the different indices are documented in the supplementary material.

## Discussion

Ethanol is an important stimulus in the environment of adult and larval *D. melanogaster* (reviewed in ^51^,^52^). The concentration of ethanol in the natural habitat of this species vary between 0.6% in ripe hanging fruits and up to 4.5% in rotting ones ^53^. In some man-made environments, such as wine cellars, the ethanol concentration can reach more than 10% ^53–55^. Given that adult *D. melanogaster* often carry yeast to the egg laying sites and inoculate ripening fruits, the induced fermentation and associated ethanol production is likely to result in higher concentrations at the larval stage ^56^. Therefore, the naturally occurring range of ethanol concentrations experienced by a larva is in agreement with most of the concentrations tested in our behavioral analysis (Fig. 1, Fig.3, and Fig. 5). Only the highest concentration tested (20% and 30% ethanol) do not occur in the natural habitat of the larva. Of course, one has to keep in mind that our experiments were performed under laboratory conditions using a substrate of agarose mixed with ethanol. Due to the evaporation of ethanol, which is indeed volatile, especially higher ethanol concentrations could have been lower in reality. However, these effects are likely to be small, as studies have shown that within the first few hours, ethanol levels drop only slightly in a 13% ethanol containing Petri dish ^55^. This corresponds approximately to the period of preparation of the plates and the conduction of the ethanol attraction and learning experiments. Similarly, evaporation seems to be of little relevance in the survival experiments, where the experimental vials were changed every day. It was shown that the ethanol concentration in vials closed with polyurethane bungs remains very stable over several days (in bottles with 6% ethanol in the medium the fall is to 3.5% after six days) ^55^.

### The ethanol attraction is based on various sensory modalities

We show that *D. melanogaster* larvae preferentially migrate to substrates containing ethanol, regardless of whether they have been previously exposed to ethanol or not. Larval attraction appears to be doses-dependent reaching a plateau between 4% and 10% (Fig.1 and Fig. 2). Our results are in agreement with several published findings that have shown that larvae prefer 1% ^57^, 2.5% ^58^, 5% ^58^, 6% ^31–33^, 10% ^57^,^58^, 17% ^34^, 20% ^58^ and even pure ethanol ^57^. Thus, our standardized experimental design provides a robust behavioral assay that allows to identify the neural and molecular basis of larval ethanol substrate choice. However, it must be mentioned that there are some studies on larval ethanol substrate choice, where the tested animals show no or only weak attraction to ethanol concentrations from 2% to 6% ^38^,^39^,^59^ and even aversion at higher concentrations ^39^,^59^. However, it is not clear whether, in addition to methodological matters such as a low volume of applied ethanol ^38^, other factors such as the enzymatic activity of Adh, the surrounding temperature, specific pre-treatments and the precise experimental procedure are responsible for this ^32–34^,^39^,^58^. Our results also suggest that certain mutations may alter the attraction to ethanol, as *w^1118^* mutants showed a reduced attraction to higher ethanol concentrations compared to the wild-type (Fig. 1).

Substrate attraction tests (Fig. 1 and Fig. 2) allow larvae to come into direct contact with ethanol. Therefore, larvae can potentially be guided by different inputs: the sense of smell or taste, but also by a caloric gain or a pharmacological effect. Examples for such cases are reported for adult *D. melanogaster* (reviewed in ^26^,^51^,^60^). Therefore, we have tried to separate some of these inputs from each other. (Please note that the pharmacological effect can only be analyzed by behavioral changes and measurement of endogenous ethanol concentration in the larvae, which were not done in our study). To investigate whether larvae specifically perceive and behaviorally respond to ethanol as an odorant, we have filled the stimulus in containers that allow ethanol to evaporate, but prevent the larvae from directly contacting it and thus excludes feeding or ingesting it (Fig. 4). This has not been studied so far. Larval ethanol attraction for the odorant stimulation was clearly evident (Fig. 4), but reduced compared to the substrate attraction of ethanol (Fig. 1). The behavioral response is almost halved from a median of 0.61 (Fig. 1) to 0.33 (Fig. 4). This means that (1) larvae indeed do perceive ethanol via the olfactory signaling pathway and (2) larvae can perceive ethanol not only as an odor. Both have to be accurately identified in the future with a standardized behavioral assay which is now available.

### Ethanol is both nutritious and toxic for the larva

Survival experiments, as we have shown in Fig. 3, are often used to analyze a positive life-prolonging effect (e.g. based on additional calories of sugar in a minimal diet), or to identify harmful effects that lead to an increased mortality of larvae (e.g. by adding bitter compounds to the food) ^18–20^. Our results suggest that ethanol performs two opposing functions. Below 8% the positive – most likely nutritional effect – increases larval survival on pure agarose; at high concentrations above 8% ethanol toxicity prevails (Fig. 3). Many early studies focused on the effects of ethanol on larval survival and increases were referred to as tolerance (reviewed in ^27^). Nowadays tolerance usually stands for an individual’s resistance to the intoxicating effects of ethanol, according to its importance for humans ^27^. Similar to our results (Fig. 3), previous work showed that ethanol consumption of up to 4% leads to increased larval fitness ^47^,^48^,^61^,^62^. Parson and colleagues showed that ethanol concentrations of 0.5%–3% allow larvae to develop to 3^rd^ instar larvae. At a concentration of 6% or higher this was not the case ^48^. Quite similar, McKechnie and Geer showed that wild-type *Canton-S* larvae transferred to ethanol concentrations of up to 4.5% partially survived into pupae ^47^. Thus our data in line with published results suggest that *Drosophila* larvae may benefit from metabolizing ethanol. Since ethanol is one of the primary alcohols, it is degraded to aldehydes which then act as acetyl-CoA in metabolic processes such as the Krebs cycle (reviewed in ^43^). Thus, ethanol contributes to the formation of adenosine triphosphate (ATP) which can be used directly or indirectly as an energy source. Studies in larvae have additionally shown that ethanol’s carbons serve as backbones for triacylglycerols and phospholipids ^61^,^63^. Consequently, it can be assumed that larvae benefit from metabolizing the quantities of ethanol that they usually encounter in their natural environment.

At higher ethanol concentrations the Adh dependent detoxification reaches its limits. In most studies, reduced survival rates occur at concentrations above 9% or 16% of ethanol ^28^,^46^,^48^ (but see also for lower concentrations ^47^). Our results are therefore comparable with most of the published work, as we show that larvae do not display increased mortality at 8% ethanol, but many of them die quickly at an ethanol concentration of 20% (Fig. 3). It was shown that high intracellular concentrations of ethanol limits the capacities to synthesis and store glycogen, lipid and protein in the larval fat body cells. Even structural changes within cells of the endoplasmatic reticulum and mitochondria were visible ^61^. Therefore, higher doses of ethanol have a deleterious effect at both the cellular and organism levels.

Surprisingly, larvae still seek out these deleterious concentration and process it as rewarding in learning experiments (Fig. 1, Fig. 3, Fig. 5 and Fig. 7). Thus, it is tempting to speculate that the answer is based on the organization of the larval microhabitats. Ethanol occurs most often as a gradient in food sources. Therefore, it might be advantageous for the larva to seek out even high ethanol sources in the first step to then consume lower substrate concentrations on site in the second step. In general, however, it appears that larvae respond differently to a given concentration of a chemosensory stimulus depending on their behavioral response. 0.3 M sodium chloride is usually repellent to larvae, but can be used as a positive teaching signal in olfactory learning experiments ^42^. 2M arabinose, a sugar that is sweet but offers no nutritional benefit, is harmful for the larva, but also highly attractive and triggers a reward function during learning ^20^. It appears that this is a general function for many chemosensory components, which may be based on mutual neuronal and genetic mechanisms.

**Figure 7:**
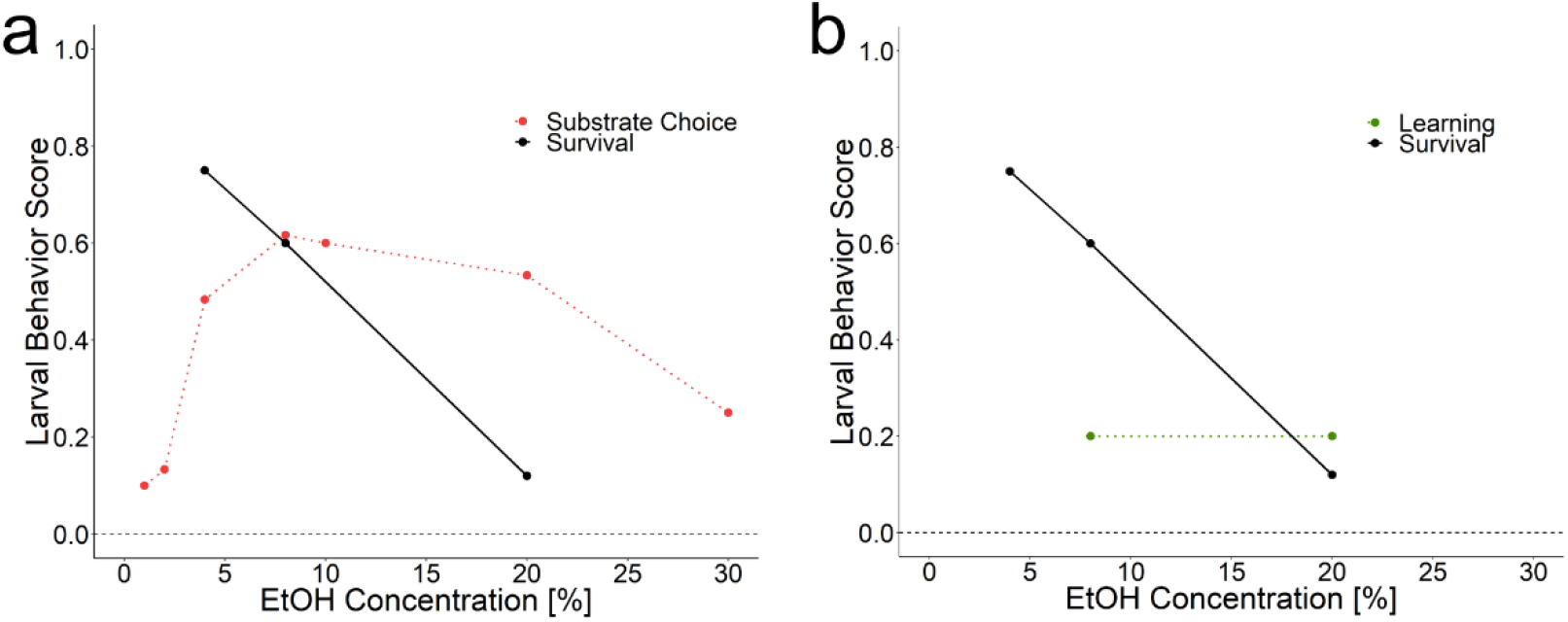
Summarizing semi schematic illustration of larval ethanol guided behavior scores. The x-axis shows ethanol concentrations from 0% to 30%. The y-axis shows the larval behavior score from 0 to 1 for *Canton-S*. Note that the data plotted here are only comparable for the corresponding ethanol concentration. (a) Substrate choice ranges from 0.1 to 0.6 (data Fig. 1b). Survival scores indicates the survival probability at day eight (data Fig. 3b). Although higher concentrations of about 20% ethanol are harmful to the larvae a preference is still detectable. The behavioral curve for ethanol seems to be shifted towards higher concentrations. (b) Ethanol dependent memory indicates the performance scores for 8 and 20% ethanol (Fig. 5a). Survival scores indicate the survival probability at day eight (data Fig. 3b). Although there is a clear negative effect on larval survival for higher concentrations of ethanol its rewarding effect stays the same.

### Ethanol provides a positive teaching signal

To our knowledge it has not yet been tested whether larvae can utilize ethanol as a positive teaching signal (unconditioned stimulus) in learning and memory assays. Our results suggest that this is the case for concentrations of 8% and 20% ethanol (Fig. 5). Therefore, not only adult flies can use ethanol as a teaching signal ^64^,^65^, but also larvae. The initial memory of adult flies, however, is aversive and only changes after 24 hours into a positive long-term memory that lasts several days ^64^. The adult behavioral response is therefore clearly different from the immediately positive larval response. Nonetheless, we gain increasing insight to understand what stimulation the larva classifies as rewarding. Chemosensory stimuli include ethanol, low salt concentrations, amino acids, ribonucleosides and various sugars (e.g., sucrose, glucose, maltodextrin, sorbitol, ribose), even though some cannot be metabolized (arabinose, xylose) ^20,21,42,49,66,67^.

We also do not yet have any insights into the neuronal and molecular organization of larval ethanol learning and memory. However, the here described assay now allows their analysis. It is worth noting that ethanol memory cannot be retrieved on a fructose test plate (Fig. 5). According to Schleyer and colleagues *D. melanogaster* larvae search for food after the conditioning phase based on their acquired experience. Thus, the odorant paired with ethanol predicts a certain gain. At the moment of testing, the animals expect such a gain and compare it with their current environmental input. It seems that the larvae do not only expect a positive gain in this situation but also its specific quality. This was shown as sugar memory could be recalled on a positive aspartic acid containing test plate and vice versa ^68^. Following the same logic, the absence of ethanol memory recall on a fructose plate thus means that larvae do not distinguish between ethanol and fructose quality. Therefore, it is possible that fructose and ethanol processing circuits overlap in the larval brain.

### Outlook

Which cells and molecules might be involved in the perception of ethanol in larvae? Based on studies on olfactory ethanol sensing in adult flies, the octopaminergic system and different olfactory receptors can be considered as an analysis entry point ^69,70^. Using the Flywalk^71^ or a two odor vial assay^72^ to measure naive adult olfactory output, it was shown that the olfactory co-receptor Orco plays an essential role in both assays when ethanol was applied. For the Flywalk analysis the effect could even be refined to the olfactory receptor genes Or42b and Or59b ^70^. At the larval stage, only Or42b is expressed in a single ORN of the dorsal organ, the main larval olfactory sense organ ^15^. Or42b responds to ethyl acetate, ethyl butyrate, propyl acetate and pentyl acetate; chemicals that are all highly attractive to larvae ^16^,^17^. It is tempting to expect that Or42b serves a similar function at the larval stage. However, ethanol has not yet been tested, and we lack any molecular and neuronal information on its perception in the larva. With respect to learning and memory, we have recently obtained a cellular understanding of the involved neuronal pathways of the larval brain. Four dopaminergic neurons of the primary protocerebral anterior medial (pPAM) cluster encode fructose dependent teaching signals and these neurons are directly connected to the mushroom body, the larval memory center ^9^,^73–76^. Hence, it is now possible to test whether aspects of the ethanol reward learning and memory are also processed by the same neuronal mushroom body network. This would also reveal a conservation of the functional patterns throughout development, since adult appetitive ethanol memory also requires dopaminergic cells and the mushroom body ^64^.

## Methods

### Fly strains

Fly strains were kept on standardized cornmeal medium containing 1% ethanol at 25°C and 65% humidity under a 14:10 h light:dark cycle. Adult flies were transferred to new food vials every 72 h. Larvae were taken from food vials and briefly washed in tap water to remove food residues. Two fly strains wild-type^CS^ and mutant *w^1118^* were used in the experiments. For experiments at specific time points during larval development, larvae with defined age were used. If not otherwise mentioned larval of different age were used. For staging larvae, adult flies were allowed to lay eggs for six hours; larvae were then collected in a 12 h cycle starting 36 h after egg laying (AEL).

### Substrate choice

For gustatory preference tests, Petri dishes (85 mm diameter; Greiner) were filled with a thin layer of 2.5% agarose substrate (VWR Life Science; type number: 97062-250). After cooling, agarose was removed from half of the plate and re-filled with 2.5% agarose substrate containing different concentration of 99.8% ethanol (1%, 2%, 4%, 8%, 10%, 20% or 30% ethanol; CHEMSOLUTE^®^; type number: 2273.1000). To keep an evaporation effect as low as possible, groups of 30 larvae of different ages were placed immediately after plate preparation (in the order of maximum 45 minutes) in the middle of the Petri dish and allowed to crawl for 5 min (or up to 120 min) at room temperature (RT) ^55^. Then, larvae were counted on each side of the Petri dish. A preference index (*Pref*) was calculated by subtracting the number of larvae on the pure agarose side (#nS) from the number of larvae on the side with a stimuli (#S), divided by the total number of larvae (#total): *Pref* = (#S – #nS) / #total. A positive *Pref* indicates attraction and a negative aversion.

### Olfactory attraction

To test for olfactory attraction, Petri dishes (85 mm diameter; Greiner) were filled with a thin layer of 2.5% agarose substrate or with 2.5% agarose substrate with different concentration of ethanol (5%, 8%, 10% and 20%). To test the olfactory stimuli of ethanol, either 8% ethanol, benzaldehyde (BA; Sigma-Aldrich, type number: 102213897; undiluted), amyl acetate (AM; Sigma-Aldrich, type number: 102172386); diluted 1:250 in paraffin oil), 1-octanol (1-Oct; Sigma-Aldrich, type number: 101858766 undiluted) or distilled water was filled into custom-made Teflon containers (4.5mm diameter) with perforated lids and placed on each side of the plate. Immediately after plate preparation, groups of 30 larvae were placed in the middle of the Petri dish and allowed to crawl for 5 min at RT. Then, larvae were counted on each side of the Petri dish. A preference index (*Pref*) was calculated by subtracting the number of larvae on the side without an odor (#nO) from the number of larvae on the side with an odor (#O), divided by the total number of larvae (#total): *Pref* = (#O – #nO) / #total. A positive *Pref* indicates an attractiveness, a negative *Pref* represents an avoidance.

### Olfactory ethanol learning and memory

All experiments were performed on Petri dishes filled with a thin layer of either 2.5% pure agarose or 2.5% agarose containing different concentrations of ethanol (8% or 20%), 0.01M fructose (Carl Roth^®^; type number: 4981.2) or 1.5M sodium chloride (Carl Roth^®^; type number: 3957.2). As olfactory stimuli, we filled 10μl AM, and 10μl BA, in custom-made Teflon containers (4.5mm diameter) with perforated lids. Immediately after plate and container preparation, groups of 30 larvae were placed in the middle of the Petri dish containing a pure agarose substrate. All experiments were conducted at RT. Larvae were exposed to AM and allowed to crawl for 5 min. Then, larvae were transferred to a new Petri dish containing a positive (ethanol, fructose) or negative (NaCl) reinforcer and exposed to BA for 5 min. After three training cycles, larvae were placed on a new Petri dish containing pure agarose substrate or agarose substrate containing 8% ethanol, 0.01M fructose, or 1.5M sodium chloride and exposed to AM and BA on opposite sides for 5 min. Then, larvae were counted on each side of the Petri dish. A second group of larvae was trained via a reciprocal training regime. For each group an independent olfactory preference index was calculated as described above. A performance index (*PI*) was calculated by adding the Pref of the first training group (*Pref1*) to the Pref of the second training group (*Pref2*) and dividing them by the number of experimental groups (#2): *PI* = (*Pref1* + *Pref2*) / #2. A positive PI indicates an attractiveness; a negative PI represents an avoidance.

### Larval survival on ethanol

To investigate larval survival in the presence of ethanol, 12 wild-type L2 stage larvae were placed in vials containing either 1% pure agarose substrate or 1% agarose substrate plus different concentrations of ethanol of 4%, 8% or 20%. Three drops of tap water were daily added to prevent larvae from dehydrating. Larvae that were alive and later the pupal stage were counted from day 1 to day 9. The percentage of survival and pupation were calculated as follows: Percentage = (number of living larvae or pupae / total number of larvae / pupae) x 100. For each condition 16 independent experimental groups were analyzed (n= 16).

### Data analysis and visualization

To test whether single groups are significantly different from zero, we used a one-sample t-test. Further statistical analyses were conducted using distribution-specific tests. For normally distributed data, we used either the paired T-test with equal variance or the one-way ANOVA followed by a TukeyHSD multiple comparison test for normal distributed data. For non-normal distributed data, we used Wilcoxon signed rank test or Kruskal-Wallis test followed by Dunn’s multiple comparison test. All statistical analyses and data visualization were done with R (V 3.5.1) in RStudio by using *stats*, *dunn.test* and *ggplot2* package. Figure panels were edited with Adobe Illustrator CS5 (San Jose, CA, USA). The significance level of statistical tests was set to 0.05; the shown confidence interval (ci) is the 95% confidence interval of the data plots. Data are displayed as box plots with the median as the middle line, the box boundaries as 25% and 75% quantiles and the whiskers as 1.5 times the interquartile range. Outliers are shown as dots directly above or below the box plots. Further details are documented in the Supplemental Data files.

## Acknowledgments

The authors thank Wolf Huetteroth, Tilman Triphan, Bert Klagges and Astrid Rohwedder for fruitful discussions and comments on the paper. This work was supported by grants from the German Research Foundation (DFG) to AST (TH1584/3-1, TH1584/6-1 and TH1584/7-1) and HS (HS675/10-1).

## Author contributions

IS and AST outlined the manuscript. IS, MB, HS and AST developed the design of the methodology; IS, MB, NN, YS and JS conducted the experiments; IS prepared and created the analysis and visualization of the data; IS, MB, HS and AST wrote and finalized the manuscript.

